# Competence-associated peptide BriC alters fatty acid biosynthesis in *Streptococcus pneumoniae*

**DOI:** 10.1101/2021.02.17.431746

**Authors:** Surya D. Aggarwal, Jessica M. Gullett, Tara Fedder, J. Pedro F. Safi, Charles O. Rock, N. Luisa Hiller

## Abstract

Membrane lipid homeostasis is required for bacteria to survive in a spectrum of host environments. This homeostasis is achieved by regulation of fatty acid chain length and of the ratio of saturated to unsaturated fatty acids. In the pathogen *Streptococcus pneumoniae*, fatty acid biosynthesis is encoded by a cluster of fatty acid biosynthesis (*fab*) genes (FASII locus) whose expression is controlled by the FabT repressor. Encoded immediately downstream of the FASII locus is BriC, a competence-induced, cell-cell communication peptide that promotes biofilm development as well as nasopharyngeal colonization in a murine model of pneumococcal carriage. Here, we demonstrate that *briC* is co-transcribed with genes of the *fab* gene cluster and that a reduction of *briC* levels, caused by decoupling its transcription from *fab* gene cluster, negatively impacts biofilm development. BriC elevates *fabT* transcription, which is predicted to alter the balance of saturated and unsaturated fatty acids produced by the pathway. We find that *briC* inactivation results in a decreased production of unsaturated fatty acids that impact the membrane properties by decreasing the abundance of di-unsaturated phosphatidylglycerol molecular species. We propose that the link between BriC, FabT and phospholipid composition contributes to the ability of *S. pneumoniae* to alter membrane homeostasis in response to the production of a quorum-sensing peptide.

**IMPORTANCE:** Adaptation of bacteria to their host environment is a key component of colonization and pathogenesis. As an essential component of bacterial membranes, fatty acid composition contributes to host adaptation. Similarly, so does cell-cell communication, which serves as a mechanism for population levels responses. While much is known about the pathways that control the biosynthesis of fatty acids, many questions remain regarding regulation of these pathways and consequently the factors that impacts the balance between saturated and unsaturated fatty acids. We find that BriC, a cell-cell communication peptide implicated in biofilm regulation and colonization, is both influenced by a fatty acid biosynthesis pathway and impacts this same pathway. This study identified a link between cell-cell communication, fatty acid composition, and biofilms and, in doing so, suggests that these pathways are integrated into the networks that control pneumococcal colonization and host adaptation.

## INTRODUCTION

*Streptococcus pneumoniae* (pneumococcus) is a major human pathogen. Worldwide, it is responsible for over one million annual deaths in children and the elderly (1). Drug resistant pneumococcus is classified as a serious threat by the CDC, such that there is a need for new therapies. Fatty acid synthesis is a core function of the cell, and as such a potential drug target.

In bacteria, fatty acids can be acquired by two independent pathways: *de novo* production or uptake from host cells. *De novo* phospholipid synthesis in pneumococcus is carried out by thirteen dissociated fatty acid synthesis genes that are a part of the FASII system. These genes are encoded in a single cluster on the genome and act to elongate and modify acetyl-CoA primers to produce saturated and unsaturated acyl chains attached to an acyl carrier protein (ACP). Attachment to ACP allows binding of any acyl chain to FASII enzymes. Uptake from the host is mediated by the fatty acid kinase (Fak) system that incorporates exogenous fatty acids into the phospholipid membrane (2, 3).

In pneumococcus, the regulation of the FASII locus is under control of FabT, which autoregulates itself and represses most FASII genes except for FabM (4, 5). FabT is constitutively expressed and binds with low affinity to the promoter regions of *fabT* and *fabK*, thereby permitting some transcription of genes in the FASII locus. Long chained acyl-ACPs, from FASII or exogenous sources, regulate FASII through FabT.

Specifically, the FabT-acyl-ACP complex which strengthens FabT’s affinity for DNA and blocks FASII gene transcription (2, 3, 6).

In the current model, the balance between saturated (SFA) and unsaturated (UFA) fatty acids requires an isomerase, FabM. FabM catalyzes the conversion of *trans*-2-to *cis*-3-enoyl-ACP (4, 5, 7). The SFA:UFA ratio is determined by the competition of FabM and FabK for the available enoyl-ACP. If FabK utilizes the enoyl-ACP, SFA are produced, and if FabM utilizes the intermediate, UFA are produced. Because FabM catalyzes an equilibrium reaction, its overexpression has little impact on UFA levels compared to the larger effects of independently manipulating either the FabK or FabF levels (Lu and Rock, 2006). Because FabT regulates FabK and FabF, but not FabM, modulation of FabT repression levels by the combination of *fabT* expression and/or acyl-ACP levels impacts the balance between SFA and UFA synthesis.

In many Gram-positive bacteria, including pneumococcus, uptake of fatty acids blocks *de novo* synthesis by triggering an inhibition of endogenous fatty acid synthesis (via FabT in the pneumococcus). For example, like *S. pneumoniae*, *Enterococcus faecalis* encodes two *acp* genes: *acpA* encoded within the fatty acid synthesis (*fab*) operon and *acpB* encoded in an operon with the acyl-ACP:phosphate transacylase *plsX*. Long chain acyl-ACP-dependent repression via exogenous fatty acids is selective for AcpB in *E. faecalis*. The transcription of two ACPs, present in different neighborhoods, ensures that acyl-ACPs originating from a host will regulate FASII synthesis; incoming acyl chains are paired with AcpB while *acpA* and *fabT* are repressed (8). In this manner many bacteria, via their ability to synthesize membrane from fatty acids acquired from the host, can survive without *de novo* synthesis as long as external sources are available. Another factor that regulates FASII is the WalRK histidine kinase signal transduction system (also known as YycFG and VicRK). Overexpression of the response regulator, WalR, modifies the expression of twelve FAS genes and results in cells phenotypically similar to *fabT* mutants that have longer-chained fatty acids (9).

Immediately downstream of the pneumococcal FASII locus is the small peptide, BriC (<u>b</u>iofilm <u>r</u>egulator <u>i</u>nduced by <u>c</u>ompetence). BriC is a ribosomally-synthesized peptide that belongs to the class of double-glycine secreted peptides in pneumococcus (10). The expression of *briC* is induced directly by ComE, the master regulator of competence, and BriC is secreted via the competence-associated ABC transporter, ComAB (11). Some pneumococcal isolates, including those from the clinically important PMEN1 and PMEN14 lineages, encode a RUPB1-containing *briC* promoter which provides a competence-independent induction of *briC* in an otherwise competence-dependent pathway. The production and secretion of BriC promote late stage biofilm development *in vitro*, and nasopharyngeal colonization in a murine model of pneumococcal carriage (11).

Here, we show that *briC* is co-transcribed with genes of the *fab* gene cluster, and that its expression modulates the membrane fatty acid composition of *S. pneumoniae*. In accordance with the role of BriC in promoting biofilm development, decreasing levels of *briC* by decoupling its transcription from the *fab* gene cluster negatively influences biofilm development. The *briC* knockout strains have altered levels of *fabT* expression coupled with a distinct shift in membrane phospholipid molecular species composition. Thus, BriC contributes to the regulation of *S. pneumoniae* FASII either directly or indirectly by altering the transcription of the FabT regulon.

## RESULTS

### *briC* is co-transcribed with genes of the *fab* gene cluster

BriC is a competence-induced gene product, yet a basal level of transcription is observed even in the absence of competence (11). The coding region for *briC* is immediately downstream of the *fab* gene cluster (159 base pairs downstream of *accA* (*spd_0390*) in strain R6D) **(Fig. 1A).** Thus, we hypothesized that the basal levels of *briC* may be attributed to its co-transcription with genes of the *fab* gene cluster. Since the genes spanning from *fabK* through *accA* are transcribed as a polycistronic unit (4), we tested whether *briC* is transcribed with the last two genes of this operon: *accD* and *accA*. We performed PCR on cDNA synthesized using RNA from planktonic pneumococcal cultures as a template and used it to determine whether transcripts extend from *accD* or *accA* to *briC* **(Fig. 1B.i)**. The results indicated that *briC* is co-transcribed with genes of the *fab* gene cluster. An *in silico* search reveals two putative promoter sequences, which may drive the co-transcription of *briC* and genes of the *fab* gene cluster (**Fig. 1A)**. The first is upstream of *fabK*, and the second is within the coding sequence of *fabG*. These promoters contain putative −35 and −10 regions, and are in agreement with promoters previously identified by Cappable-seq (12).

**Fig. 1.**
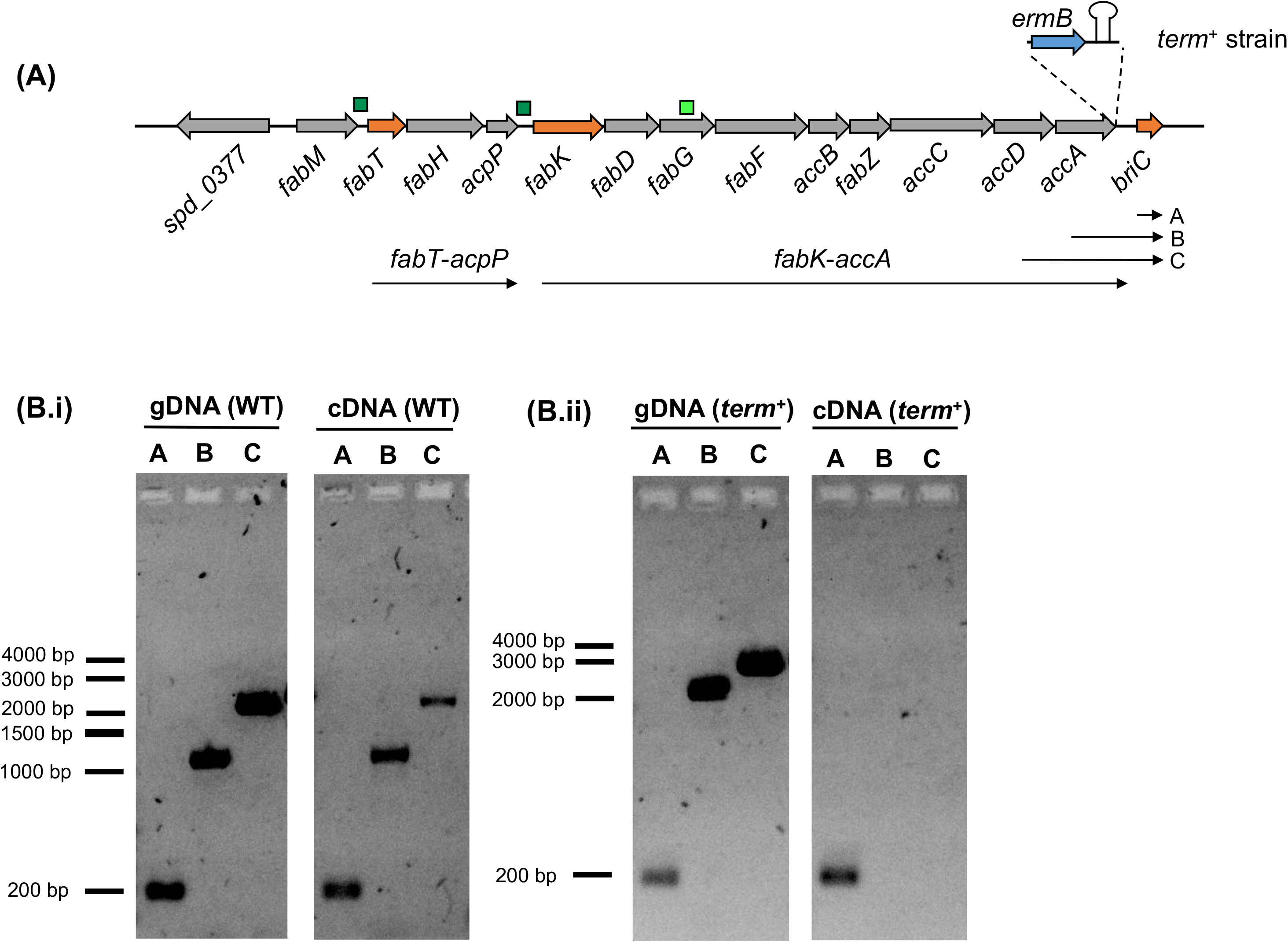
*briC* is co-transcribed with genes of the *fab* gene cluster. **(A)** Genomic organization of the *fab* gene cluster and *briC*. The *fab* gene cluster consists of thirteen genes ranging from *fabM* to *accA*. FabT regulates the expression of two operons: *fabT*-*acpP* and *fabK*-*accA*. *briC* is situated downstream of *accA*. Small dark green boxes indicate promoters with a FabT-binding site, while light green box indicates an additional putative promoter. Inset: *term*^+^ strain contains *ermB* cassette followed by the terminator B1002 immediately downstream of *accA*. Transcripts *fabT*-*acpP* and *fabK*-*accA* are labelled. Labels A, B and C refer to the different transcripts tested below. **(B.i)***briC* is transcriptionally linked to genes of the *fab* gene cluster. gDNA and cDNA from WT strain were amplified using primers expected to produce three different amplicons: A: *briC* only (177bp), B: *accA*-*briC* (1088bp), C: *accD*-*briC* (1912bp), visualized on agarose gel. **(B.ii)** Insertion of the terminator relieves co-transcription of *briC* with *accA*. gDNA and cDNA from *term*^+^ strain was amplified using primers expected to produce amplicons A, B and C (*as above*), visualized on agarose gel. cDNA for *term*^+^ shows a positive band only for *briC* alone. Amplicons B & C from gDNA have a higher molecular size in *term*^+^ relative to WT because of the presence of *ermB* and terminator (additional 1051bp).

To study the importance of co-regulation of *briC* with genes of the *fab* gene cluster, we opted to decouple the transcription of *briC* from that of the *fab* gene cluster. We generated a strain with a transcriptional terminator immediately downstream of *accA* (*term*^+^ strain). As expected, while a transcript with *briC* alone is present in the *term*^+^ strain, the *accA-briC* transcript is no longer detected (**Fig. 1B.ii**). Thus, introduction of the terminator relieved *briC* of its co-transcription with genes of the *fab* gene cluster. Further, the competence-dependent induction of *briC* was preserved in the *term*^+^ strain (*briC* was induced 3.26-fold following CSP treatment). We conclude that *briC* expression can be regulated in concurrence with the *fab* gene cluster, as well as independently via CSP.

### Co-expression of *briC* with genes from the *fab* gene cluster contributes to biofilm development

We have previously demonstrated that BriC promotes biofilm development (11). Since *briC* can be co-transcribed with the upstream fatty acid genes, we hypothesized that decoupling *briC* from fatty acid synthesis would negatively impact biofilm development. In support, we observe an approximately 15% reduction in biofilm biomass and thickness in the *term*^+^ cells relative to WT cells, when testing biofilms at 72h post-seeding on abiotic surfaces (**Fig. 2A, B**). While maximum thickness is a measure of the distance of highest point or the peak from the bottom layer containing biomass, the average thickness over biomass is an indicator of the general shape and spatial size of the biofilm.

**Fig. 2.**
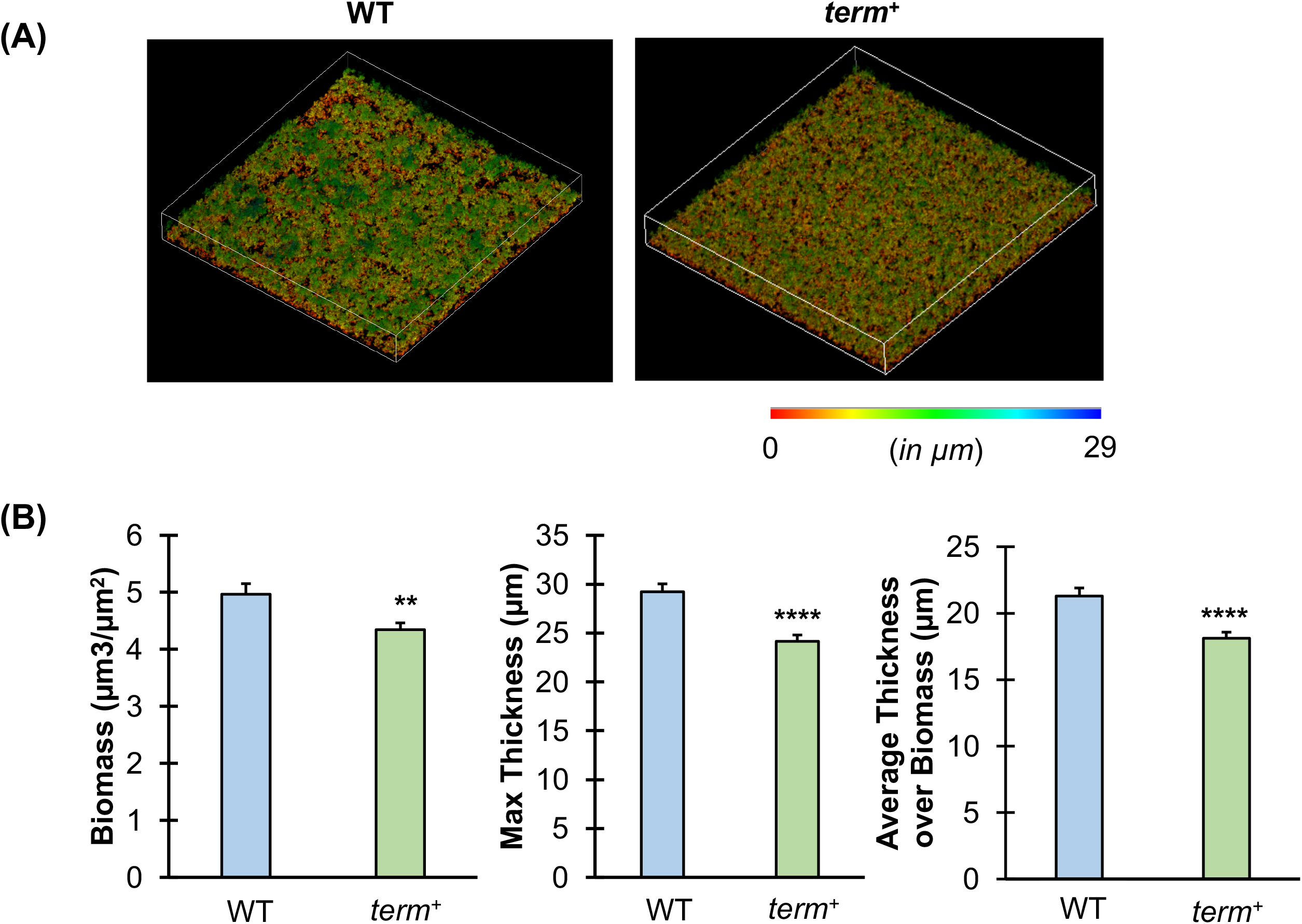
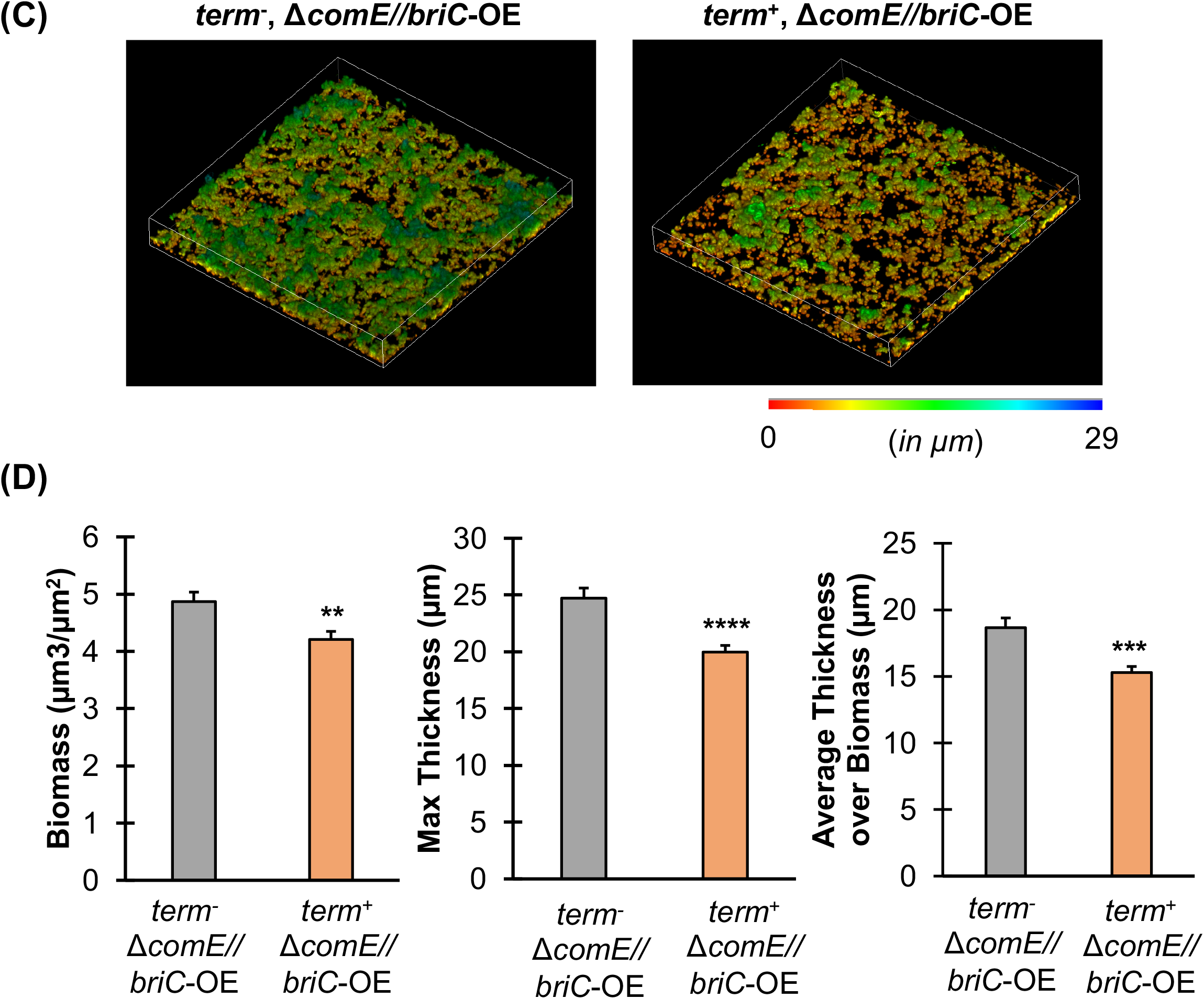
Co-expression of *briC* with the *fab* gene cluster promotes biofilm development. **(A, C)** Representative confocal microscopy images showing top view of the reconstructed biofilm stacks of *term*- and *term*^+^ cells in **(A)** WT and **(C)**Δ*comE*//*briC*-OE genomic background of strain R6D stained with SYTO59 dye at 72h. Images are pseudo-colored according to depth (scales shown). **(B, D)** COMSTAT2 quantification of 72h biofilm images. Y-axis denotes units of measurement: μm^3^/μm^2^ for biomass, and μm for maximum thickness and average thickness over biomass. Error bars represent standard error of the mean calculated for biological replicates *(n = 3)*; ** *p*<0.01, *** *p*<0.001 and **** *p*<0.0001 using Student’s *t*-test.

To establish whether the biofilm defect was associated with an alteration in the competence-dependent induction of *briC*, we tested biofilm development in cells where *briC* was regulated in a competence-independent fashion. We made use of a strain where *briC* is overexpressed due to a promoter that encodes a RUP sequence (P*briC*_long_-*briC*), and where *comE* is deleted (henceforth, referred to as Δ*comE*//*briC*-OE). We have previously shown that expression of *briC* from P*briC*_long_ bypasses the impact of *comE* deletion on biofilm development (11). Thus, we tested whether decoupling transcription of *briC* from the *fab* gene cluster by the introduction of terminator (*term*^+^) influences biofilms in the Δ*comE*//*briC*-OE background. Akin to the WT background, presence of the terminator (*term*^+^) leads to a significant reduction in biomass and thickness of biofilms in the Δ*comE*//*briC*-OE strain compared to the *term*^−^ strain in the same background (**Fig. 2C, D**). These results strongly suggest that induction of *briC* via control of the *fab* gene cluster contributes to the role of BriC in promoting biofilm development. Thus, regardless of the mechanism of induction, increased expression of *briC* positively contributes to biofilm development.

### BriC contributes to membrane compositional homeostasis

BriC is a secreted peptide that is co-transcribed with genes of the *fab* gene cluster. Owing to this genomic organization, we investigated whether BriC played any functional role in altering fatty acid synthesis in pneumococcal cells. FabT regulates genes of the *fab* gene cluster, including itself (4). To test the impact of BriC on expression of *fabT*, we generated a fusion of the *fabT* promoter with *lacZ* and measured the β-galactosidase activity in WT and Δ*briC* strains. We observed approximately a 35% decrease in β-galactosidase activity in Δ*briC* relative to WT cells **(Fig. 3)**. We conclude that BriC enhances *fabT* transcription from its autoregulated promoter. The reduced expression of *fabT* signals repression of the entire regulon suggesting that the absence of *briC* expression would alter the membrane phospholipid composition. Specifically, the repression of the FabT regulon would increase UFA biosynthesis at the expense of SFA (4).

**Fig. 3.**
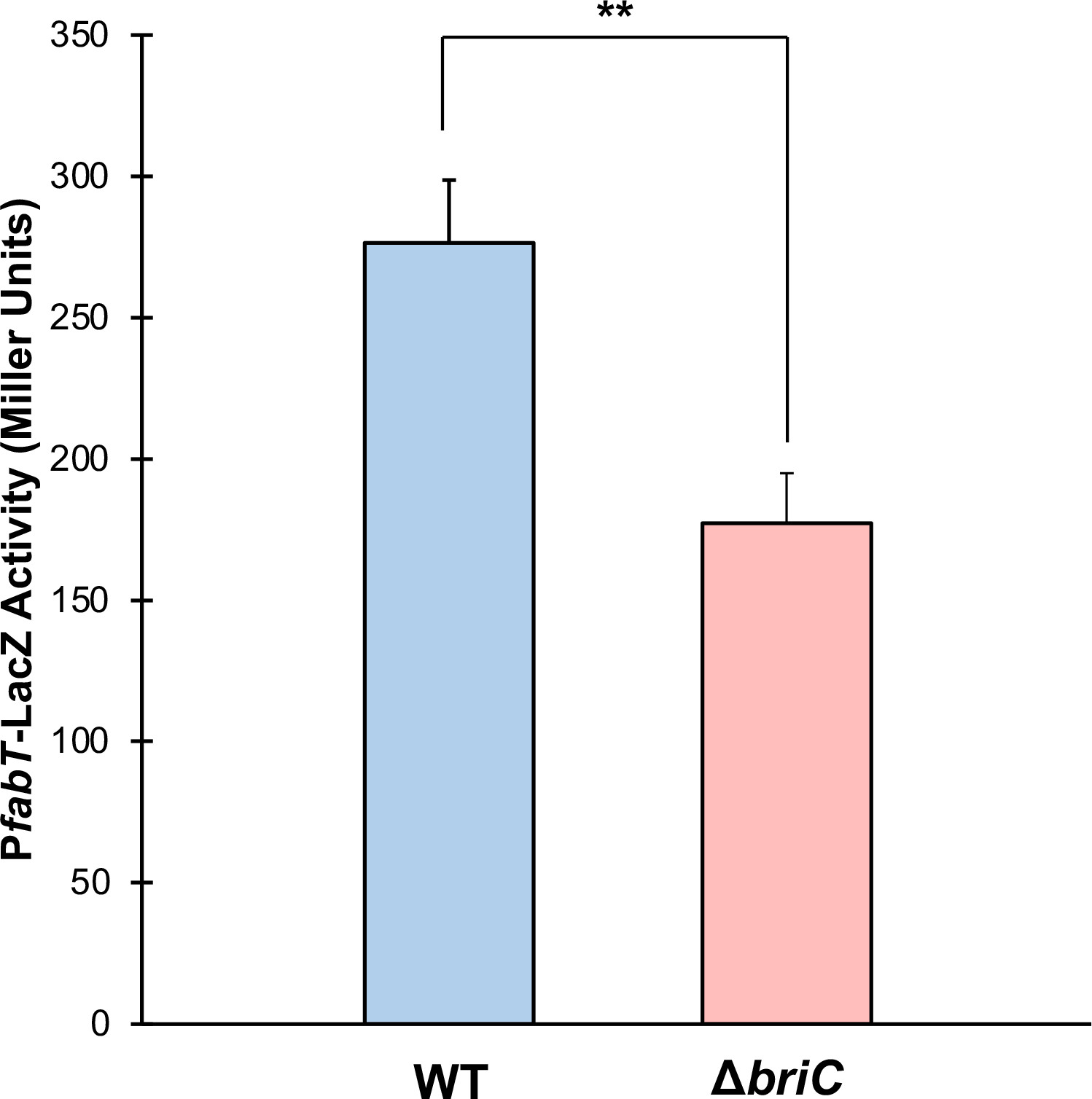
BriC induces the levels of *fabT*. β-galactosidase assay comparing the LacZ activity of *fabT* promoter in WT and Δ *briC* cells. Cells were grown in TY medium until mid-log phase. Y-axis denotes promoter activity in Miller Units expressed in nmol p-nitrophenol/min/ml. Error bars represent standard error of the mean for biological replicates *(n = 3)*; “ns” denotes statistically non-significant comparison, ** *p*<0.01 using Student’s *t*-test.

This prediction was tested by determining the composition of the membrane phospholipids in wild-type and Δ*briC* strains. Pneumococcus uses the FASII system to produce acyl chains which are transferred, via positionally-specific acyltransferases, to the 1- and 2-positions of glycerol-3-phosphate (G3P) that determine the composition of phosphatidic acid, the precursor to all membrane glycerolipids. We employed liquid chromatography-mass spectrometry (LC-MS) to determine the phosphatidylglycerol (PG) membrane molecular species composition in wild-type, Δ*briC*, and *briC*-overexpressing cells (*briC*-OE) cells. LC-MS analysis determines the total carbon number of acyl chains in the 1- and 2-position of the G3P backbone, as well as the number of double bonds, and is an accurate representation of the acyl chain production of pneumococcal strains.

The wild-type strain made primarily mono- and di-unsaturated PG molecular species with the predominant peaks containing 32, 34, or 36 carbons (**Fig. 4A**). The molecular species distribution in these samples consisted of 16:0, 16:1, 18:0, and 18:1 acyl chains and is typical of other pneumococcal strains (2, 13). The acyl chains comprising the predominant peaks correspond to 16:0/16:1 (32 carbons), 18:1Δ11/16:1 and 16:0/18:1Δ11 (34 carbons), and 18:1Δ11/18:1Δ11 (36 carbons) (2). In the Δ*briC* strain, we observed a reduction in the unsaturated molecular species at each carbon number in comparison to the wild-type strain (**Fig. 4B**). None of these changes were observed in the *briC*-OE cells relative to the WT strain (**Fig. 4C**). The quantification of three replicates shows a consistent decrease in the unsaturated PG molecular species at each carbon number (**Fig. 4D**). We conclude that FASII of the Δ*briC* strain produces a lower amount of unsaturated fatty acids that, in turn, alters the membrane phospholipid molecular species composition. Together, these results suggest that BriC, via *fabT*, alters the lipid composition of pneumococcal cell membranes, tilting the balance toward unsaturated fatty acids.

**Figure 4.**
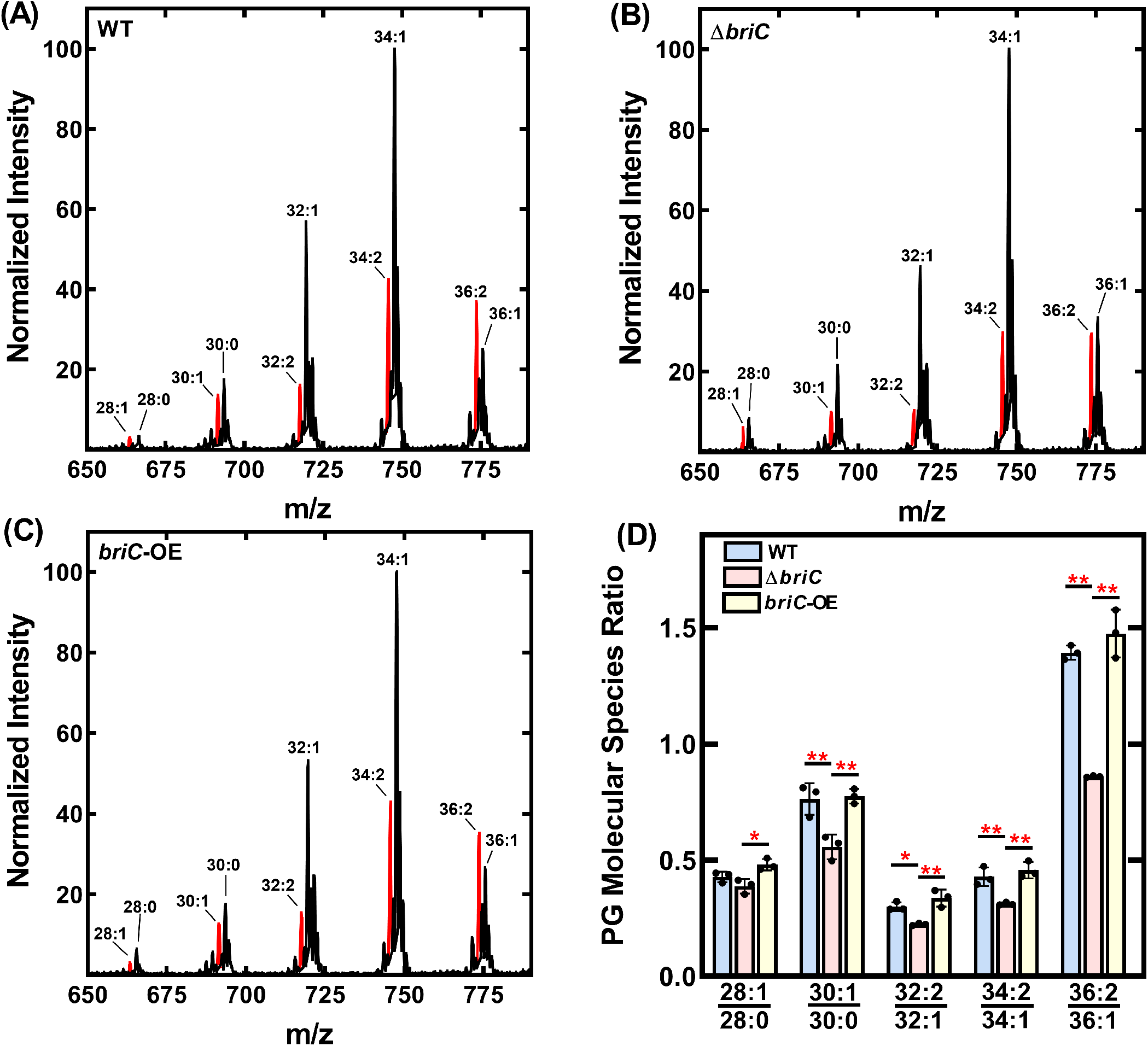
BriC promotes a membrane composition enriched in unsaturated fatty acids. Mass spectrometry analysis of the PG molecular species of R6D pneumococcal strain and its isogenic mutants. Representative spectrum of PG molecular species of **(A)** WT, **(B)** Δ*briC*, and **(C)** *briC*-OE strains. The most unsaturated molecular species for each carbon number are highlighted in red. **(D)** The PG species from three biological replicates were quantified and the ratios for each carbon number were calculated. *p-* values were calculated in Prism software using ANOVA followed by Tukey’s test for multiple comparisons.

## DISCUSSION

Long chain fatty acids serve as essential components of bacterial membranes and compositional changes therein are essential for cellular survival and environmental adaptation. The biosynthesis of fatty acids is tightly regulated. While much is known about the regulation and synthesis of the cell membrane, gaps remain regarding the nature of the molecular signals that activate this pathway. Here we present evidence that BriC, a small, secreted peptide implicated in cell-cell communication, is co-transcribed with genes of the *fab* gene cluster and participates in regulation of phospholipid membrane composition via a role in *fabT* induction. Moreover, as we have previously shown that BriC promotes biofilms, we demonstrate that this phenotype is linked to co-regulation between FASII genes and *briC.*

The demonstration that *briC* is co-transcribed with the *fab* gene cluster, reveals the third regulatory pathway for *briC* regulation. The regulation of *briC* by ComE, via the ComE-binding box, is conserved across strains in the species and demonstrates a tight link between competence induction and *briC* expression (11). In a subset of strains, the transcription of *briC* is also influenced by the presence of transposable RUP (**<u>r</u>**epeat **<u>u</u>**nit of **<u>p</u>**neumococcus) sequence in its promoter (11). RUP allows for a CSP-independent pathway for the expression of *briC*. The discovery that *briC* can be co-transcribed with the *fab* gene cluster, reveals a third pathway for regulation. Further, it suggests that BriC may be regulated by multiple two component systems, ComE and WalRK. WalRK triggers the activation of the *fab* gene cluster and *briC* is co-transcribed with genes of this locus, thus it is likely that WalRK promotes *briC* expression. The activation of *briC* by multiple pathways is consistent with a gene network where BriC is positioned to respond to diverse regulatory inputs. Signaling through WalRK is important in maintenance of cell shape, division, pathogenesis and in the response to stresses such as oxidative stress (14–16). Competence activation has also been described of as a general SOS response pathway, as well as a sensor of cell density. Thus, the colonization factor BriC may respond to cell density and stress conditions and induce changes in membrane composition and biofilm growth.

We have previously demonstrated that BriC promotes biofilm development and nasopharyngeal colonization (11). In this study, we show that BriC influences membrane lipid composition. Are these phenotypes connected? A study comparing transcriptional profiles of pneumococcal cells growing in biofilm versus planktonic mode of growth found an upregulation of fatty acid biosynthesis genes during biofilm development (17). A role for fatty acid biosynthesis and metabolism in biofilm formation has also been reported in other bacteria including *Bacillus subtilis*, *Staphylococcus aureus* and *Pseudomonas aeruginosa* (18–21). It seems plausible that BriC-dependent changes in membrane properties contribute to biofilm development. Alternatively, enhanced cell-cell signaling associated with a biofilm-mode of growth, may enhance BriC-mediated effects on lipid composition, and serve as a link between biofilms and lipid composition.

In this work, we have revealed a link between a cell-cell communication peptide and the regulation of fatty acid composition. Our findings reveal that *briC* is co-regulated with the *fab* gene cluster, and, reciprocally, that it impacts membrane homeostasis by influencing transcription of the FabT regulon.

## MATERIALS & METHODS

### Bacterial strains & growth conditions

The experimental work was performed with the R6D wild-type strain of *Streptococcus pneumoniae* (Hun663.tr4) as this was used in our previous studies of BriC (11). Colonies were grown from frozen stocks by streaking on TSA-II agar plates supplemented with 5% sheep blood (BD BBL, New Jersey, USA). Unless otherwise stated, streaked colonies were picked and inoculated in fresh Columbia broth (Remel Microbiology Products, Thermo Fisher Scientific, USA) whose pH was adjusted to 6.6 by the addition of 1M HCl and thereafter, incubated at 37°C and 5% CO2 without shaking.

### Construction of mutants

Mutant strains were constructed by using site-directed homologous recombination and selected by the addition of an antibiotic resistance marker. The *term*^+^ transformation construct was generated by ligating the amplified flanking regions with antibiotic resistance cassette followed by transcriptional terminator B1002. Between 1-2kb of flanking regions upstream and downstream of the region of interest were amplified from parental strain using Q5 2x Master Mix (New England Biolabs, USA). The antibiotic resistance gene *ermB* was amplified from *S. pneumoniae* SV35-T23. The sequence of the terminator was added to the primers. The PCR products were assembled by Gibson assembly using NEBuilder HiFi DNA Assembly Cloning Kit (New England Biolabs, USA).

### Bacterial transformations

Bacterial target strains were grown in acidic Columbia broth until an OD_600_ of 0.05 and followed by addition of 125μg/mL of CSP1 (sequence: EMRLSKFFRDFILQRKK; purchased from GenScript, NJ, USA) and 1μg of transforming DNA. The cultures were incubated at 37°C and 5% CO2 without shaking for 2 hours followed by plating on Columbia agar plates containing the appropriate antibiotic: kanamycin (150μg/ml), erythromycin (2μg/ml) and incubating overnight. Resistant colonies were cultured in selective media, and the colonies confirmed using PCR. Bacterial strains generated in this study are listed in Supplementary Table S1.

### RNA extractions

For qRT-PCR, samples were grown until an OD_600_ of 0.1, followed by CSP1 treatment for 0 and 10 minutes. This was followed by addition of RNALater to preserve RNA quality and pelleting of cells. The cells were lysed by resuspending the pellet in an enzyme cocktail (2mg/ml proteinase K, 10mg/ml lysozyme, and 20μg/ml mutanolysin). Then, RNA was isolated using the RNeasy kit (Genesee Scientific, USA) following manufacturer’s instructions. Contaminant DNA was removed by treating with DNase (2U/μL) at 37 °C for at least 45 mins followed by RNA purification using the RNeasy kit. The RNA concentration was measured by NanoDrop 2000c spectrophotometer (Thermo Fisher Scientific, USA). The purity of the RNA samples was confirmed by the absence of a DNA band on an agarose gel obtained upon running PCR products for the samples amplified for *gapdh*.

### qRT-PCR

Purified RNA was used as a template for first-strand cDNA synthesis by using qScript cDNA Synthesis Kit (Quantabio, USA) followed by qRT-PCR using PerfeCTa SYBR Green SuperMix (Quantabio, USA) in an Applied Biosystems 7300 Instrument (Applied Biosystems, USA).16S rRNA counts were used for normalization.

### Biofilm development assay

For biofilm development assays, pneumococcal cells were grown in acidic Columbia broth until the cultures reached an OD_600_ of 0.05. Then, 3mls of culture was seeded on 35MM glass bottom culture dishes (MatTek Corporation, USA) and incubated at 37°C and 5% CO2 without shaking. At 24h and 48h post-seeding, the supernatant from the dishes was carefully aspirated with a pipette, followed by the addition of the same volume of pre-warmed media made at one-fifth of the original concentration. The biofilms were fixed for analysis at 72h post-seeding. For fixation, supernatants were aspirated, and the biofilms were washed thrice with PBS to remove non-adherent and weakly adherent cells. Thereafter, biofilms were fixed with 4% paraformaldehyde (Electron Microscopy Sciences, USA) for 20 minutes. The biofilms were then washed with PBS three times and stained for confocal microscopy.

### Confocal microscopy & quantification of biofilms

SYTO59 Nucleic Acid Stain (Life Technologies, USA) was used to stain biofilms as per manufacturer’s instructions for 30 minutes. The stained biofilms were then washed three times and preserved in PBS for imaging. Imaging was performed on the stage of Carl Zeiss LSM-880 META FCS confocal microscope, using 561nm laser for SYTO dye. Z-Stacks were captures at every 0.46 μm, imaged from the bottom to the top of the stack until cells were visible, and reconstructed in Carl Zeiss black edition and ImageJ. The biofilm stacks were analyzed using COMSTAT2 plug-in for ImageJ (22) and the different biofilm parameters (biomass, maximum thickness, and average thickness over biomass) were quantified. For depiction of representative reconstructed Z-stacks, empty slices were added to the images so the total number of slices across all the samples were the same. The reconstructed stacks were pseudo-colored according to depth using Carl Zeiss black edition.

### Construction of *lacZ* fusions

Chromosomal transcriptional *lacZ*-fusions to the target promoters were constructed as previously described (11). Briefly, *lacZ*-fusions were generated in the *bgaA* gene using modified integration plasmid pPP2. The *fabT* and *fabK* promoter regions were amplified from R6D strains and modified to contain KpnI and XbaI restriction sites. The products were then digested with restriction enzymes followed by sticky-end ligation of the products. These plasmids were transformed into *E*. *coli* TOP10 strain, and selected on LB (Miller’s modification, Alfa Aesar, USA) plates, supplemented with ampicillin (100μg/ml). The plasmids were then purified by using E.Z.N.A. Plasmid DNA Mini Kit II (OMEGA bio-tek, USA), and transformed into pneumococcal strains and selected on Columbia agar plates supplemented with kanamycin (150μg/ml).

### β-galactosidase assay

β-galactosidase assay was performed as previously described (23). For assaying the β-galactosidase activity, cells were grown in TY media (TH medium supplemented with 0.5% yeast extract) until exponential phase and frozen. The frozen cells were thawed and re-inoculated in TY media and grown until mid-exponential phase for analysis.

### Membrane Lipid Composition Analysis

Bacterial cells were inoculated in CDM-Glucose and incubated at 37°C and 5% CO2 without shaking until they reached an OD_600_ of 0.5. CDM-Glucose was prepared as previously described (PMID: 23505518). The cells were then pelleted by centrifuging at 4000rpm for 15 minutes followed by washing with PBS three times. The PBS was decanted and the washed cells were frozen at −20°C before being resuspended in 1 ml deionized water and vortexed. Lipids were resuspended in chloroform:methanol (2:1) and extracted using the Bligh and Dyer method (24). PG was analyzed using a Shimadzu Prominence UFLC attached to a QTrap 4500 equipped with a Turbo V ion source (Sciex). Samples were injected onto an Acquity UPLC BEH HILIC, 1.7 μm, 2.1 × 150 mm column (Waters) at 45°C with a flow rate of 0.2 ml/min. Solvent A was acetonitrile, and solvent B was 15 mM ammonium formate, pH 3. The HPLC program was the following: starting solvent mixture of 96% A / 4% B, 0 to 2 min isocratic with 4% B; 2 to 20 min linear gradient to 80% B; 20 to 23 min isocratic with 80% B; 23 to 25 min linear gradient to 4% B; 25 to 30 min isocratic with 4% B. The QTrap 4500 was operated in the Q1 negative mode. The ion source parameters for Q1 were: ion spray voltage, −4500 V; curtain gas, 25 psi; temperature, 350°C; ion source gas 1, 40 psi; ion source gas 2, 60 psi; and declustering potential, −40 V. The system was controlled by the Analyst® software (Sciex). The sum of the areas under each peak in the mass spectra was calculated, and the percent of each molecular species present was calculated with LipidView software (Sciex).

### Statistical tests

For comparisons between only two groups, student’s *t*-test was performed. *p*-values of less than 0.05 were considered to be statistically significant. Statistical analyses of the ratios of PG molecular species were determined using an ANOVA and Tukey’s Test.

## Acknowledgements

We thank Matthew Frank for mass spectrometry and Jason Rosch for thoughtful suggestions. This research was supported by NIGMS (GM034496 to CR), the American Lebanese Associated Charities (CR), NIAID (R01 AI139077-01A1 to NLH), the Eberly Family Trust (NLH) and Glen de Vries Fellowship (SDA). The content is solely the responsibility of the authors and does not necessarily represent the official views of the National Institutes of Health.

---

**Table S1:** Strains used in this experimental work.

| Strain ID | Strain Name | Description | Source |
| --- | --- | --- | --- |
| LH339 | R6D | Penicillin-resistant R6-derivative, Hun663.tr4 | Severin *et al.* 1996 |
| LH65 | R6D $\Delta briC$ | *briC* (*spr_0388*) replaced with *ermB* in R6D; Ery<sup>R</sup> | Aggarwal *et al.* 2018 |
| LH825 | *term*<sup>+</sup> | *ermB* ligated with terminator inserted downstream of *accA* in R6D | This study |
| LH366 | $\Delta comE//briC$-OE | *comE* (*spr_2041*) replaced with *aad9* in R6D *PbriC*<sub>long</sub><sup>-</sup> *briC*; Spec<sup>R</sup>, Kan<sup>R</sup> | Aggarwal *et al.* 2018 |
| LH845 | *term*<sup>+</sup>, $\Delta comE//briC$-OE | *ermB* ligated with terminator inserted downstream of *accA* in $\Delta comE::PbriC$<sub>long</sub>-*briC*; Spec<sup>R</sup>, Kan<sup>R</sup>, Ery<sup>R</sup> | This study |
| LH920 | *E. coli* TOP10 (*PfabK-lacZ*) | *E. coli* strain transformed with modified pPP2 containing R6D *fabK* promoter fused with *lacZ*; Kan<sup>R</sup>, Amp<sup>R</sup> | This Study |
| LH921 | *E. coli* TOP10 (*PfabT-lacZ*) | *E. coli* strain transformed with modified pPP2 containing R6D *fabT* promoter fused with *lacZ*; Kan<sup>R</sup>, Amp<sup>R</sup> | This Study |
| LH922 | R6D (*PfabK-lacZ*) | R6D *fabK* promoter fused with *lacZ* in R6D; Kan<sup>R</sup> | This Study |
| LH925 | R6D (*PfabT-lacZ*) | R6D *fabT* promoter fused with *lacZ* in R6D; Kan<sup>R</sup> | This Study |
| LH928 | $\Delta briC$ (*PfabK-lacZ*) | R6D *fabK* promoter fused with *lacZ* in R6D $\Delta briC$; Ery<sup>R</sup>, Kan<sup>R</sup> | This Study |
| LH931 | $\Delta briC$ (*PfabT-lacZ*) | R6D *fabT* promoter fused with *lacZ* in R6D $\Delta briC$; Ery<sup>R</sup>, Kan<sup>R</sup> | This Study |
| LH360 | R6D::*PbriC*<sub>long</sub>-*briC* | *briC*, along with PN4595-T23 *briC* promoter inserted downstream of *bga* in R6D; Kan<sup>R</sup> | Aggarwal *et al.* 2018 |

**Table S2:**
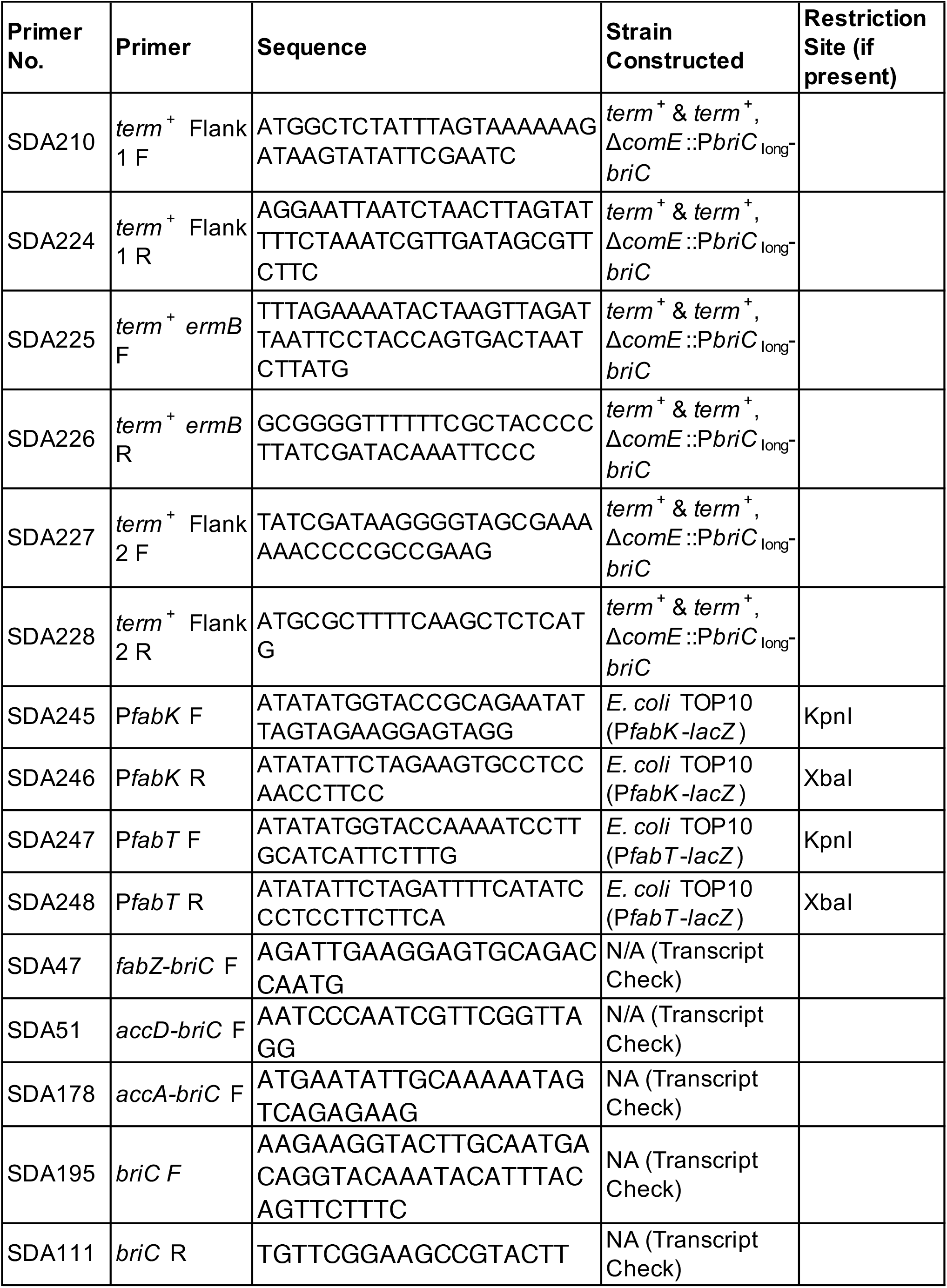
Primers used in this study.

